# Estimation of allele-specific fitness effects across human protein-coding sequences and implications for disease

**DOI:** 10.1101/441337

**Authors:** Yi-Fei Huang, Adam Siepel

## Abstract

A central challenge in human genomics is to understand the cellular, evolutionary, and clinical significance of genetic variants. Here we introduce a unified population-genetic and machine-learning model, called **L**inear **A**llele-**S**pecific **S**election **I**nferenc**E** (**LASSIE**), for estimating the fitness effects of all potential single-nucleotide variants, based on polymorphism data and predictive genomic features. We applied LASSIE to 51 high-coverage genome sequences annotated with 33 genomic features, and constructed a map of allele-specific selection coefficients across all protein-coding sequences in the human genome. We show that this map is informative about both human evolution and disease.

## Introduction

Innovations in DNA sequencing and genotyping have enabled the discovery of millions of genetic variants in human populations, with new variants continuing to be discovered at a rapid pace^1–4^. The great majority of these variants, however, are likely to have no impact on cellular function or human phenotypes, including disease, and many others are probably of only minor importance. The task of identifying which genetic variants are functionally important remains a major rate-limiting step in human genetics, with implications for both basic research and clinical practice.

Numerous computational strategies have been developed for the identification of functional variants, both to prioritize variants for experimental follow-up, and to address broader issues such as the genetic architecture of disease or the fraction of human nucleotides that are functionally important^5–10^. These computational predictors generally leverage genomic features correlated with function, such as sequence conservation^6, 8, 11–13^, protein structure^14–16^, chromatin accessibilit^y1718^, and protein-DNA interactions^19, 20^. Recently it has been shown that predictive power can be boosted by considering multiple features together, typically using supervised machine-learning models such as logistic regression, random forests, or support vector machines^21–25^. These models detect complex patterns associated with known pathogenic variants and use them to predict the effects of unannotated variants, often with good accuracy.

Nevertheless, the existing supervised machine-learning predictors suffer from some important limitations. For example, their predictions are typically hard to interpret, because they reflect some measure of similarity to a training set of known pathogenic variants based on a complex statistical model rather than a model formulated in terms of biological principles. In addition, the “known” disease variants used for training are generally unrepresentative of all pathogenic variants—e.g., by being enriched for coding regions, splice sites, and well-studied genes^23 26—^ which results in training biases and poor generalization. A related problem is that the reported prediction power for these methods is typically over-optimistic, because it is based on held-out training data with the same biases^26^. In general, these methods effectively serve as predictors of variants *like those in the training set*, rather than of all functional variants of interest.

An alternative strategy is to identify genetic variants that are subject to purifying (negative) selection. This approach depends on the assumption that functional and disease-associated variants are likely to reduce evolutionary fitness, which clearly does not hold in all cases. Nevertheless, this approach has the important advantages of mitigating the bias from training data and allowing for more interpretable, evolution-based models. This evolution-based strategy has now been used explicitly or implicitly by many state-of-the-art variant prioritization methods, including **LINSIGHT**, fitCons, CADD, and FunSeq2^9, 27–32^. Among these methods, **LINSIGHT** and fitCons are based on explicit evolutionary models and can be used to obtain maximum-likelihood estimates of interpretable quantities, such as the per-nucleotide probability that new mutations will have fitness consequences. These methods perform well in the prioritization of disease and regulatory variants and also provide evolutionary insights^9, 27^, but they have some important limitations. For example, **LINSIGHT** and fitCons assume that all alternative alleles at each nucleotide have equal effects on fitness^9, 27, 28, 33^, and do not provide estimates of true selection coefficients, which arguably provide the most precisely interpretable description of fitness effects.

A separate thread in the population genetics literature has addressed the problem of estimating the bulk distribution of fitness effects (DFE) from a designated collection of genomic regions, such as all coding sequences. Methods for addressing this problem typically calculate the probability of a summary of polymorphism data, such as the site frequency spectrum (SFS), given an explicit parameterization of selection coefficients using diffusion approximations of the Wright-Fisher model^34–40^. These methods generally also make use of an explicit model of demographic history, because of the confounding effect of demography on the SFs^36, 37, 41^. These DFE inference methods allow for the inference of true selection coefficients but they are unable to pinpoint the fitness effects of individual variants owing to the intrinsic sparsity of polymorphisms.

In this article, we present a unified model that combines elements of machine-learning methods for variant prediction and diffusion-approximation methods for DFE inference to enable estimation of allele-specific selection coefficients at every nucleotide in a genomic region of interest. We have implemented our model in a computer program, called **L**inear **A**llele-**S**pecific **S**election **I**nferencE (**LASSIE**), and applied it to all protein-coding sequences in the human genome, using publicly available human polymorphism data and more than two dozen predictive genomic features. We validate LASSIE by comparing the estimated selection coefficients with known patterns of natural selection in coding regions and by testing their predictive power for inherited pathogenic variants and cancer-driver mutations. We then show that LASSIE can be used to define a “null” distribution for the relationship between genomic features and selective effects, which enables the identification of genes under unusually strong or unusually weak selection. Genes under unusually strong selection are associated with brain-specific expression and autism spectrum disorder.

## Results

### LASSIE uses a unified machine-learning and population genetic model to estimate allele-specific selection coefficients

The key idea behind the LASSIE model is that, while polymorphisms are too sparse to allow direct estimation of allele-specific selection coefficients, there is a strong correlative relationship between genomic features and fitness effects that can be exploited to enable such estimation. The general idea is similar to that behind fitCons^9, 28^and LINSIGHT^27^, but in this case a richer machine-learning model accommodates allele-specific effects and a diffusion-based likelihood function allows for the estimation of true selection coefficients.

The LASSIE model consists of two components (**Fig. 1**). First, for the population genetic component of the model, we use a generative probabilistic model for the site frequency spectrum (SFS), adopting the Poisson Random Field (PRF) framework for direct likelihood calculations^34, 42, 43^. Second, we account for predictive genomic features using a neural network. The output of this network is not a class assignment, as in typical supervised-learning applications, but instead is a set of parameters that feed into the PRF model for likelihood calculations. Thus, the overall model is a generative model for the data, fitted in an unsupervised manner by maximization of the likelihood, but it conditions on a potentially large, complex, and informative set of genomic features using a neural network. This conditioning allows for pooling of data across genomic sites, and improved shrinkage estimators for allele-specific selection coefficients.

**Figure 1:**
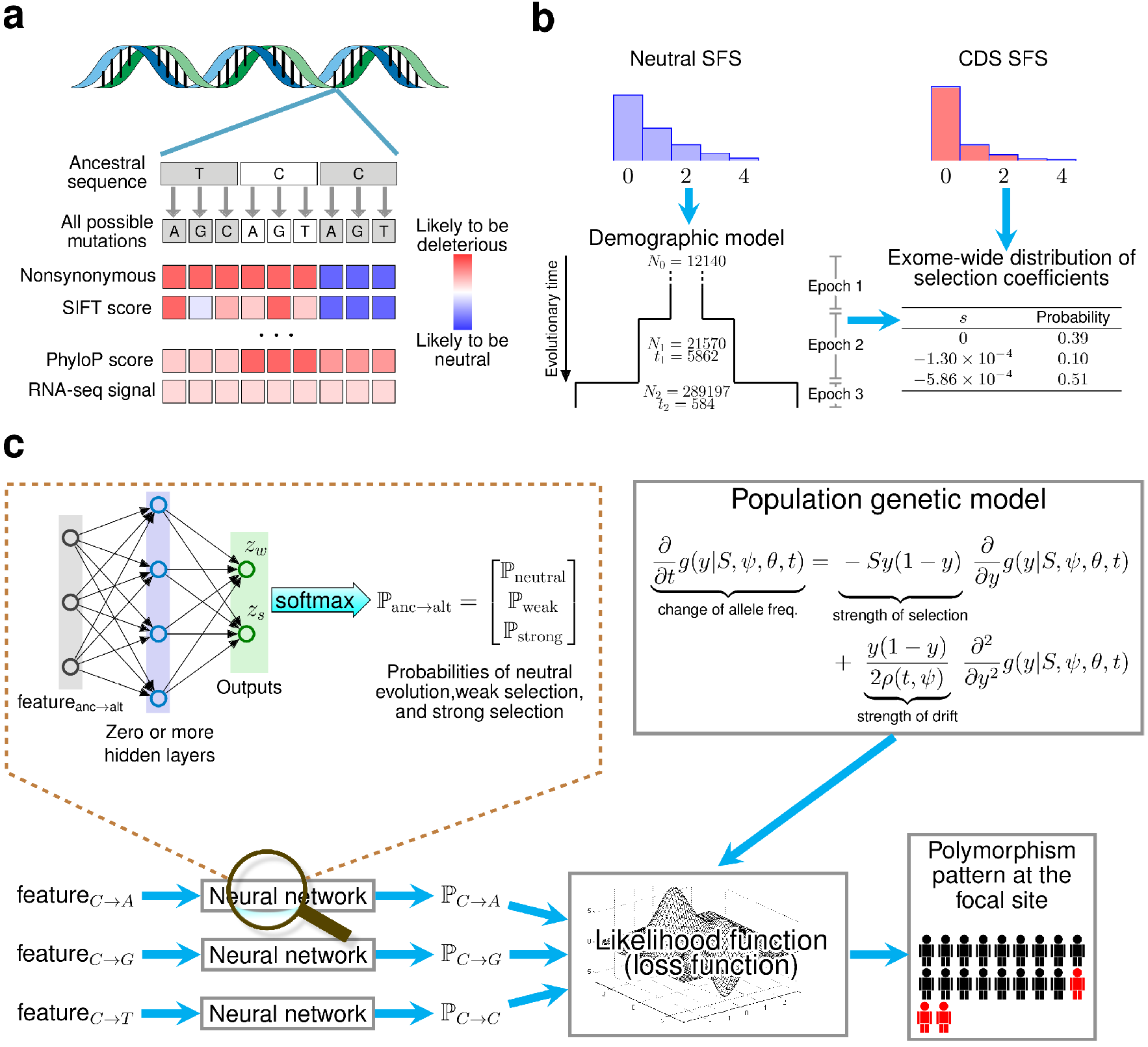
Overview of LASSIE. (**a**) For each potential protein-coding mutation, we collected 33 genomic features likely to be informative about natural selection, including variant categories, protein and nucleotide conservation scores, and RNA-seq signals (**Supplementary Table 1**). (**b**) A three-epoch demographic model was fitted to the site-frequency spectrum (SFS) for putatively neutral exon-flanking sequences for 51 high-coverage Yoruba genomes sequences. A mixture model for neutral evolution (*s* = 0), weak negative (*s* = –1.30 × 10^−4^), and strong negative (*s* = −5.86 × 10^−4^) selection was then fitted to the SFS for coding sequences (CDS; see Methods). (**c**) A mixture density network model defines the probabilities of the three components of the mixture model (ℙ_neutral_, ℙ_weak_, and ℙ_strong_) for each possible mutation at each nucleotide site, conditional on the local genomic features. These probabilities allow the likelihood of the polymorphism data to be computed under the Poisson Random Field (PRF) model, using diffusion approximation methods. The parameters of the network are estimated by maximum likelihood, by treating the (negative) log likelihood as a loss function for the neural network. After training, the weights for the three mixture components define a coarse-grained distribution of fitness effects for all potential mutations at each site. This distribution is summarized by a single expected value of |*s*| for each mutation.

### Population genomic data is described using a PRF-based mixture model

For reasons of efficiency, selection is accommodated using a three-component mixture model rather than a contin-uous distribution of selection coefficients. The mixture components capture the average effects of strong negative selection, weak negative selection, and neutral drift, respectively. The use of a mixture model allows the full PRF model to be fitted to the data in a preprocessing step, so that only the site-specific probabilities of the mixture components (the mixture coefficients) need to be estimated in the context of the neural network (see below).

To account for the confounding influence of demography on the SFS, we first fitted a simple demographic model to a collection of putatively neutrally evolving nucleotide sites flanking protein-coding exons. We focused on the 51 high-coverage Yoruba samples from the 1000 Genomes Project, because this population appears to be well described by a pure “expansion” model, without population bottlenecks or introgression events^34, 37, 39^. We assumed a three-epoch model with a constant effective population size in each epoch, and we estimated the timings and magnitudes of population expansions by maximum likelihood (Methods). The estimated model posits that the Yoruba population experienced a moderate expansion about 6000 years ago, followed by a more dramatic expansion about 600 years ago (Fig. 1b). Despite its simplicity, this demographic model provides an excellent fit to the observed SFS (Supplementary Fig. 1).

We then fitted a mixture model to genome-wide protein-coding sequence data, estimating the three mixture coefficients as well as the selection coefficients for the weak and strong negative selection components but keeping the neutral model fixed. This analysis indicates that ~10% of potential coding mutations in the human exome are under weak negative selection with a representative selection coefficient of *s* = –1.30 × 10^−4^ and about 51% of coding mutations are under strong negative selection with a representative selection coefficient of *s* = –5.86 × 10^−4^ (Fig. 1b).

We found that this mixture model also fit the exome-wide SFS well (**Supplementary Fig. 1**).

### Genomic features are incorporated using a mixture density network model

We modeled the relationship between genomic features and allele-specific probabilities of selection components using a mixture density network model^44^. This model transforms the genomic features associated with each allele to site- and allele-specific mixture coefficients (Fig. 1c). These mixture coefficients, in turn, can be used to compute the probability of the exome-wide data under the PRF mixture model. Thus, the edge weights in the neural network function as the free parameters of a generative model for population genomic data conditional on genomic features.

Hypothesizing that genomic features used in variant prioritization would also be informative in this context, we collected 33 diverse features for every potential derived mutation in the human exome, including protein conservation scores, nucleotide conservation scores, protein structural features, RNA-seq signals, and categories indicating changes in the encoded protein (nonsynonymous, nonsense, stop-gained; Fig. 1a; see **Supplementary Table 1** for a complete list of features).

We then fitted the mixture density model to the exome-wide data by maximum likelihood, keeping the selection coefficients fixed at their previously estimated values (see Methods). The features most strongly predictive of selection included stop-gain and missense mutations, several measures of phylogenetic conservation, and several features describing structural properties of amino acid substitutions such as whether the affected reside is buried or exposed, or whether the substitution is predicted to stabilize or destabilize folding^45^ (**Supplementary Fig. 2**). For ease of interpretation, we summarized the probabilities of neutrality, weak negative, and strong negative selection for each candidate mutation by the absolute value of the expected selection coefficient, |*s*|.

Notice that our general framework allows for a variety of network architectures, ranging from many-layered networks to the simple case of no hidden layers. The use of hidden layers provides the potential to capture nonlinear relationships between genomic features and selection coefficients, but at the cost of larger numbers of free parameters and increased risk of overfitting. In our tests, we found that a linear model actually fit the data better than a nonlinear one (**Supplementary Table 2**), so we have adopted this simple “linear” architecture for LASSIE.

### The estimated selection coefficients are consistent with known evolutionary patterns but suggest pervasive weak selection against synonymous mutations

The selection coefficients estimated by LASSIE are highly variable across potential mutations (see Fig. 2a & b). As expected, LASSIE assigns larger values of |*s*| to nonsynonymous and nonsense mutations than to synonymous mutations. Because synonymous mutations tend to occur at third codon positions, the spatial distribution of allele-specific selection coefficients exhibits a general three-nucleotide periodic pattern in coding regions. Inspection of individual genes reveals that LASSIE frequently distinguishes known pathogenic variants (shown in red, with relatively large estimates of |*s*|, in Fig. 2a & b) from benign variants (shown in blue, with small estimates of |*s*|).

**Figure 2:**
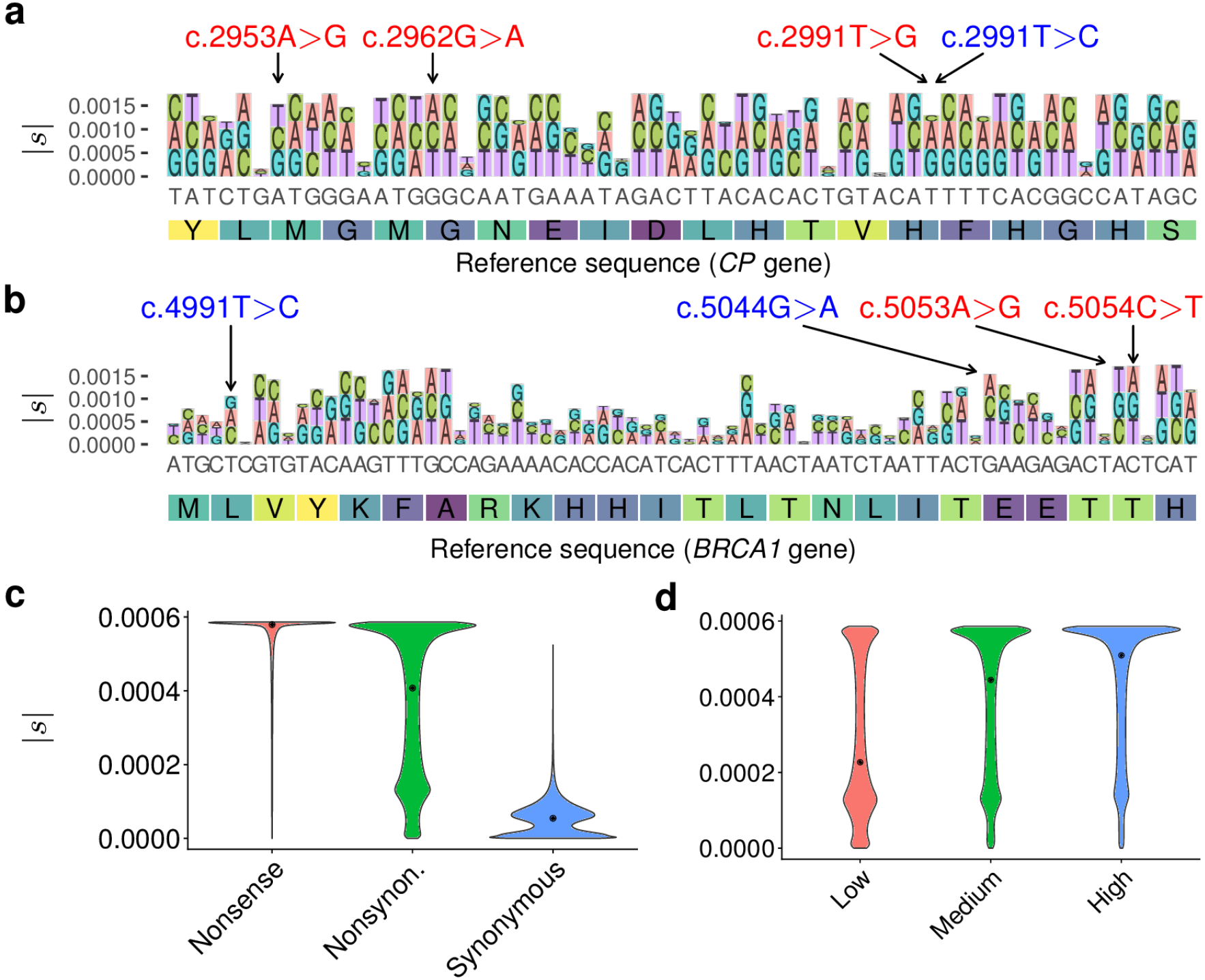
Distributions of selection coefficients estimated by LASSIE. (**a**) Variant-specific selection coefficients |*s*| estimated for all potential mutations in a 60-bp region in the *CP* gene. Nonreference alleles are distinguished by color and drawn with height proportional to |*s*|. The y-axis indicates cumulative |*s*|. The three nonsynonymous variants indicated at top in red (c.2953A>G, c.2962G>A, and c.2991T>G) are associated with Mendelian diseases and the synonymous variant indicated in blue (c.2991T>C) is benign. (**b**) Estimates of |*s*| for all potential mutations in a 72-bp region in the *BRCA1* gene. The two nonsynonymous variants indicated at top in red (c.5053A>G and c.5054C>T) are pathogenic and the two nonsynonymous variants indicated in blue (c.4991T>C and c.5044G>A) are benign. (**c**) Distributions of estimated selection coefficients |*s*| for nonsense, nonsynonymous, and synonmyous mutations. (**d**) Distributions of |*s*| for genes expressed at low, medium, and high levels, based on tertiles of RNA-seq readcounts from Roadmap Epigenomics data^82^, showing a positive correlation between expression level and |*s*| (Spearman’s rank correlation coefficient ρ = 0.338; *p* < 10^−15^, two-tailed *t*-test).

Overall, the distribution of |*s*| recapitulates well-known patterns of constraint on coding sequences. For example, LASSIE predicts that most nonsense mutations are under strong negative selection (Fig. 2c). In contrast, nonsynonymous mutations show a bimodel distribution of selection coefficients, with modes corresponding to strong and weak negative selection (Fig. 2c). While coarse-grained and truncated at our estimate for strong negative selection (see Discussion), this distribution is reasonably consistent with the bulk distribution inferred in a non-site- and allele-specific manner in previous studies; for example, we estimate that an expected 45% of nonsynonymous mutations are under neutral or weak negative selection(|*s*| ≤ 1.3 × 10^−4^ in our formulation) in comparison to estimates of ~30% with |*s*| ≤ 1.0 × 10^−4^ in ref. 37. In agreement with previous analyses based on codon substitution models^46^, we also find that nonsynonymous mutations in highly expressed genes are under significantly stronger negative selection than nonsynonymous mutations in genes expressed at lower levels (Fig. 2d).

Interestingly, the distribution of |*s*| for synynomous mutations suggests that only an expected 70.5% of such mutations are effectively neutral, whereas 25.9% are under weak negative selection and 3.6% are under strong negative selection (Fig. 2c). Weak negative selection on synonymous mutations is significantly elevated in highly expressed genes, multi-exon genes, and SRSF1 and SRSF7 binding sites (**Supplementary Fig. 3**), suggesting that roles in mRNA splicing contribute to it, perhaps among other features. This finding of a substantial influence from weak negative selection on synonymous substitutions is consistent with studies showing reduced substitution rates or reduced nucleotide diversity at synonymous sites relative to pseudogenes or introns^47–52^and suggests that the widespread practice of using such mutations as a proxy for neutral evolution^49-53^ could result in major biases in downstream analyses (see Discussion).

### The estimated selection coefficients are predictive of mutations associated with Mendelian diseases and cancer

While LASSIE was designed as an evolutionary measure, it may also be useful in the prediction of mutations associated with disease, assuming such mutations tend to be under selection^11, 12, 27, 29, 30^. To evaluate the method in this setting, we measured its power in the prediction of known Mendelian disease variants, comparing it with the popular variant prioritiza-tion methods PolyPhen-2^21^, SIFT^11^, Eigen^54^, CADD^29^, and phyloP^8^. In this experiment, we used pathogenic and benign variants from the ClinVar database^55^ as positive and negative examples, respectively. Despite no use of disease data for training (see Discussion), LASSIE performed remarkably well on this benchmark (Fig. 3a), displaying slightly greater values of the area under the receiver operating characteristic curve (AUC) statistic (AUC=0.879) than even the best previously published methods, such as Eigen (AUC=0.867) and PolyPhen-2 (AUC=0.845).

**Figure 3:**
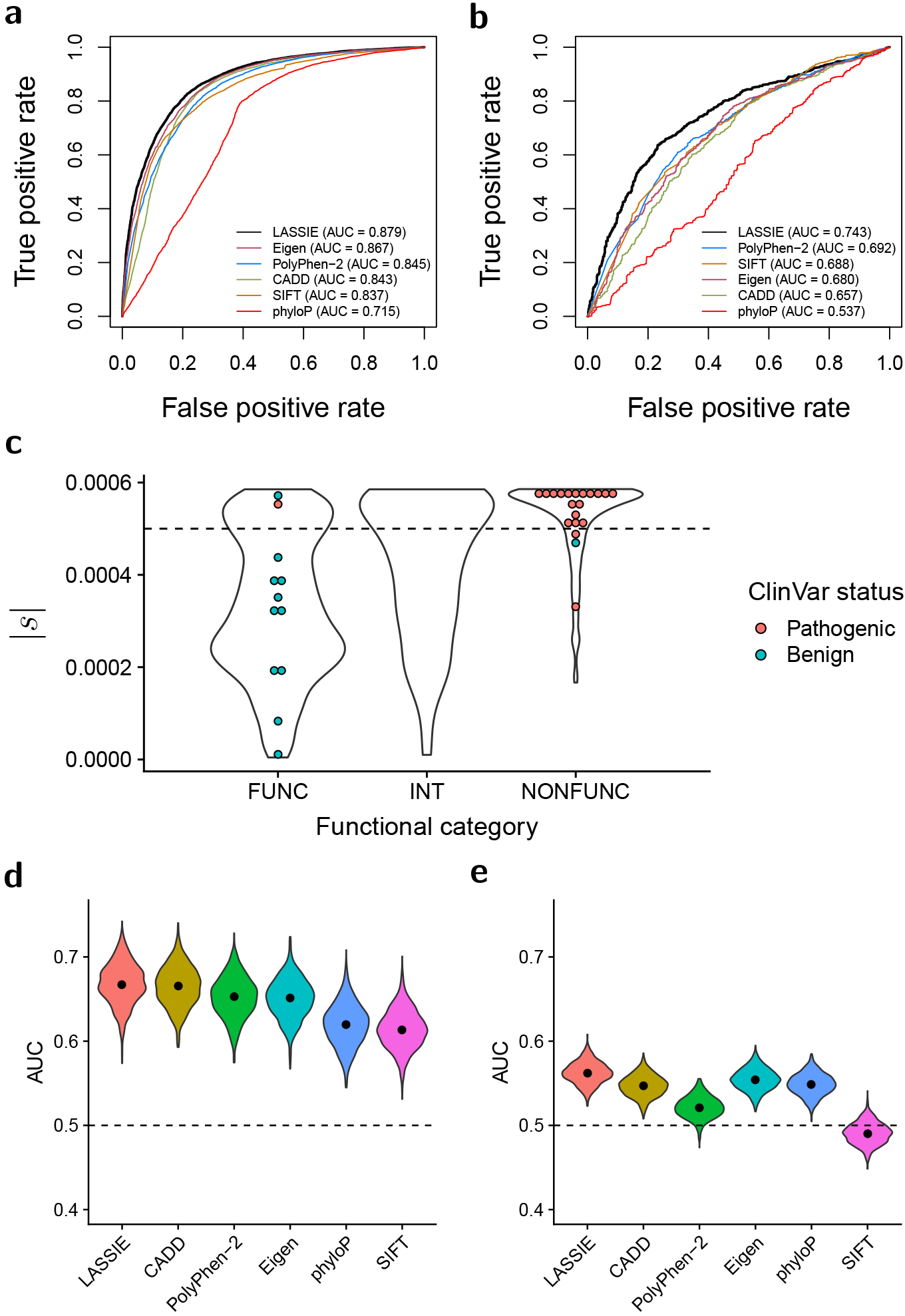
Performance in predicting disease-associated nonsynonymous variants. Performance is quantified using the area under the receiver operating characteristic curve (AUC) statistic. Results for LASSIE are compared with those for Eigen^54^, PolyPhen-2^21^, CADD^29^, **SIFT**^11^, and phyloP^8^. (**a**) Performance for pathogenic variants from ClinVar^55^. (**b**) Performance for cancer driver mutations from ref. 56. (**c**) Distributions of estimated |*s*| for variants in ***BRCA1*** predicted to be “functional” (FUNC; i.e., nondisruptive), “intermediate” (INT), or “non-functional” (NONFUNC; i.e., disruptive) by saturation genome editing^57^. Colored dots indicate those variants also having expert-reviewed status in ClinVar (CLINREVSTAT=reviewed_by_expert_panel). (**d**) Performance for rare (MAF < 1%) GWAS hits. (**e**) Performance for common (MAF > 5%) GWAS hits, showing that all methods have limited power.

As a second, largely orthogonal, test of predictive power for clinically relevant variants, we evaluated LASSIE’s performance in the prioritization of nonsynonymous cancer-driver mutations. Cancer-driver mutations in germline cells may significantly increase the risk of early-onset malignant tumors and, therefore, are likely to be under strong purifying selection in human populations. To test this hypothesis, we obtained a set of nonsynonymous mutations overlapping with mutational hotspots recurrently observed across patients in 243 cancer genes^56^, which should be enriched for cancer drivers. We randomly sampled a matched number of singleton nonsynonymous somatic mutations in the same genes to represent putative passenger mutations. LASSIE showed reasonable accuracy in this task (AUC=0.743), again performing better than all other methods tested, although the overall power was modest for all predictors (Fig. 3b). Nevertheless, the selection coefficients estimated by LASSIE are significantly predictive of cancer-driver mutations and could potentially be combined with other features to improve predictive power.

To examine disease relevance at higher resolution, we compared our estimates of |*s*| with a recent saturation genome editing (SGE) study of 13 exons of the ***BRCA1*** gene^57^. This study assigned nearly every possible single nucleotide variant (SNV) in these exons a “function score” indicating its effect on cell growth in an optimized HAP1 cell line, and then classified each SNV as “non-functional” (i.e., disruptive to growth), “functional” (nondisruptive), or “intermediate.” We found that “non-functional” variants had mostly high estimates of |*s*|, “functional” variants were enriched for medium and low estimates of |*s*|, and “intermediate” variants had intermediate estimates of |*s*| (Fig. 3c). Moreover, the 20 “pathogenic” variants from ClinVar that have been reviewed by experts almost all were both classified by SGE as “nonfunctional” and had close to the maximum possible estimate of |*s*|, whereas the 12 expert-reviewed “benign” variants from ClinVar were almost all “functional” and tended to have low to moderate estimates of |*s*| (Fig. 3c). Indeed, a threshold of |*s*| = 0.0005 (dotted line) would almost perfectly distinguish between the pathogenic and benign variants in ClinVar, with only three misclassifications (see **Supplementary Fig. 4** for ROC curves). Nevertheless, many “functional” variants appear to be under fairly strong selection, and some “non-functional” variants under fairly weak selection, indicating that there are fundamental limits to the use of natural selection as an indicator for disease (see Discussion).

### LASSIE and other methods have reasonable accuracy for rare but not common GWAS variants

Because rare genetic variants are most likely to be under negative selection, we hypothesized that evolution-based methods would be more predictive of rare than common variants associated with complex traits. To evaluate this hypothesis, we tested several methods separately on rare (MAF < 1%) and common (MAF > 5%) nonsynonymous variants from the GWAS catalog^58^, using matched variants from the 1000 Genomes Project as negative controls (Methods). In agreement with our hypothesis, most predictors were significantly more powerful in the prediction of rare GWAS variants than of common variants (Fig. 3d-e). Furthermore, LASSIE was among the most accurate methods in the prioritization of both rare and common GWAS variants.

### Brain-specific and autism spectrum disorder-related genes are under unusually strong selec-tion

LASSIE’s assumption of a single shared relationship, across all genes, between predictive genomic features and |*s*| may fail for certain subsets of genes. We searched for groups of genes that systematically deviate from this average relationship, using rare variants from 60,706 exomes in the ExAC data set to obtain high-resolution information about strong negative selection^4, 59^. To characterize the “null” distribution for the observed number of nonsynonymous variants per gene, we used a previously estimated context-dependent mutation rate map^60^ to describe site-specific mutation rates. This model predicted numbers of rare synonymous variants that were generally well correlated with the observed data (**Supplementary Fig. 5**). We then combined these site-specific mutation rates with the probabilities of strong negative selection under LASSIE to obtain an expected rate of rare nonsynonymous variants per each gene. Finally, we computed *p*-values for the observed numbers of rare (MAF < 0.001) nonsynomous variants per gene with respect to these expected rates under a Poisson-Binomial model (see Methods). We refer to the genes having significantly fewer variants than expected as being under *enhanced selection,* and the genes with significantly more variants than expected as being under *relaxed selection.*

Among 11,602 autosomal genes examined, we identified 1,118 genes and 773 genes as being under significantly enhanced or relaxed selection, respectively (Fig. 4a; FDR rate < 0.001). Interestingly, we found that the genes under enhanced (strong negative) selection were more likely to be exclusively expressed in the central nervous system (CNS) or associated with autism spectrum disorder^61, 62^(Fig. 4b). These genes were also enriched in Gene Ontology terms and pathways associated with the CNS^63^ (**Supplementary Tables 3 & 4**). The genes under relaxed selection, by contrast, tended to be exclusively expressed in liver or skeletal muscle (Fig. 4b) and to be involved in fundamental metabolic pathways (**Supplementary Tables 5 & 6**).

**Figure 4:**
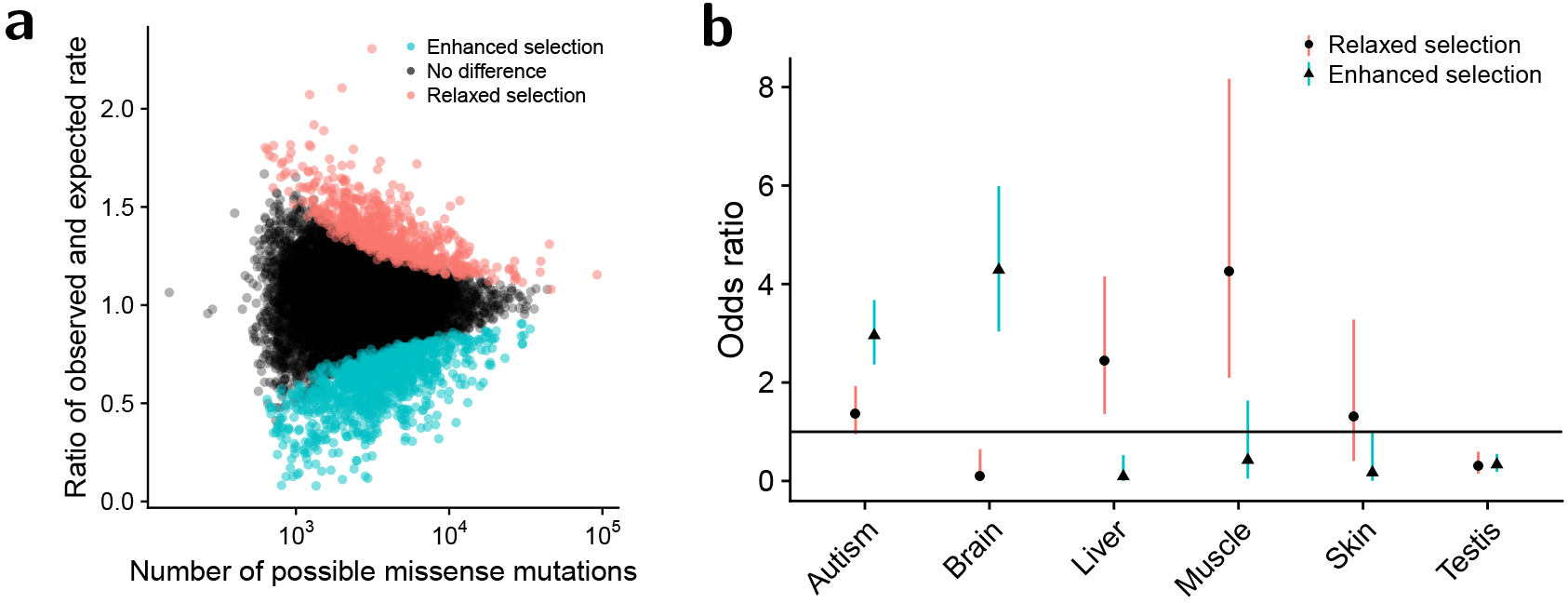
Genes under “enhanced” or “relaxed” selection relative to the exome-wide LASSIE model. (**a**) Number of potential missense mutations per gene (*x*-axis) vs. fold-change of the observed number of rare missense mutations relative to the number expected under a Poisson-Binomial null model based on LASSIE (*y*-axis; see Methods). Each dot represents a single protein-coding gene. Dots for genes showing significantly more rare variants than expected (“relaxed”; *n* = 773) are colored red, whereas those for genes showing significantly fewer rare variants than expected are colored blue (“enhanced”; *n* = 1118). (**b**) Groups of genes enriched for enhanced or relaxed selection. Dots represent odds ratios of enrichment, with bars indicating 95% confidence intervals. Genes under enhanced selection tend to be exclusively expressed in the central nervous system or associated with autism spectrum disorder. In contrast, genes under relaxed selection tend to be exclusively expressed in liver and skeletal muscle.

## Discussion

In this article, we have introduced LASSIE, the first computational method for estimating allele-specific selection coefficients at individual nucleotides across the human genome. LASSIE unifies ideas from the literature on variant prioritization and the literature on the bulk distribution of fitness effects (DFE). Like most methods for DFE inference, LASSIE is based on a generative model for allele frequencies, which can be fitted to the data by maximum likelihood without the need for labeled training data. At the same time, the LASSIE model is explicitly conditioned on a rich set of genomic features similar to those considered by variant prioritization methods. Using a flexible neural-network design, LASSIE pools polymorphism data across sites having similar genomic features to obtain improved estimates of selection coefficients. We have used LASSIE to generate a map of |*s*| for all possible single nucleotide variants in human protein-coding genes (available as a UCSC Genome Browser track: http://compgen.cshl.edu/LASSIE/), based on 51 high-coverage Yoruba genomes and 33 predictive genomic features.

For reasons of efficiency, we chose to approximate the full DFE using a mixture model, with components corresponding to neutral drift, weak negative, and strong negative selection. This strategy allows the model to be rapidly fitted to genome-wide data, but results in a rather coarse-grained estimate of the DFE. In our formulation, this approach ignores positive selection, which we have previously shown is difficult to detect in this setting^9^. Importantly, this strategy is also limited in its ability to make distinctions among large values of |*s*|, because estimates are effectively truncated at the value for the “strong” mixture component, |*s*| = 5.9 × 10^−4^. As a result, LASSIE appears to substantially underestimate |*s*| for nonsense mutations, for which the true value may be as much as two orders of magnitude larger^64^. A related problem is that our analysis is based on only 51 genome sequences, whereas more precise estimates will depend on much larger data sets (ref. 64 considered >60,000 exomes). In principle, our framework could be extended to infer full continuous distributions of s from larger data sets, but such an extension would require a number of technical improvements, including relaxation of the infinite-sites assumption underlying our model for the site frequency spectrum, accurately accounting for the effects of very recent explosive population growth^4, 65^, and further improvements to computational efficiency.

Strikingly, we find that about 30% of synonymous mutations are under negative selection. Selection on synonymous mutations appears to be almost exclusively weak, rather than strong, suggesting a limited impact on disease. Nevertheless, weak selection will tend to prohibit the fixation of synonymous alleles and will reduce synonymous substitution rates. Indeed, at a selection coefficient of *s* = –1.3 × 10^−4^ (our estimate for the corresponding mixture component), nearly all selected synonymous mutations would eventually be lost, rather than fixed, causing the observed synonymous substitution rate to be reduced by almost 30% relative to the neutral rate. This projection is consistent with a number of previous analyses of human and other mammalian data^47–52^, which have observed reductions of ~20-40% in substitution rates or nucleotide diversity at synonymous sites. Overall, it appears that negative selection on synonymous mutations is far more common than once believed, even in humans, with important implications for the widespread practice of using synonymous sites to estimate the neutral substitution rate.

The value of evolutionary methods for disease prediction ultimately depends on the degree to which natural selection correlates with disease risk. While many disease-associated variants show signatures of selection, it stands to reason that some will not, for example, because they are associated with late-onset diseases or diseases whose prevalence is strongly associated with features of modern life. Conversely, many potential variants that show signatures of selection will not relate to disease, for example, because they are strongly deleterious at embryonic or even pre-fertilization (e.g., in sperm competition) stages and never appear in patients; because they are deleterious only in the presence of a no-longer-existing genetic background; or because they reduce fitness without disrupting normal health (as through sexual selection). Nevertheless, the relationship between natural selection and Mendelian disease is sufficiently strong that evolutionary methods are fairly effective at identifying pathological variants in databases such as ClinVar, with LASSIE performing as well or better, in our experiments, than any other available computational method—including well-established methods such as PolyPhen-2 and SIFT. Interestingly, LASSIE also significantly outperformed other methods in the prioritization of nonsynonymous cancer-driver mutations, despite not being designed for the unique features of somatic evolution.

We found that recently published measures of the functional impact of point mutations in ***BRCA1*** based on saturation genome editing (SGE)^57^ correlated fairly well with LASSIE’s measure of natural selection, both across all mutations and for the subset in ClinVar (Fig. 3c). Nevertheless, an expected 44% of variants considered “functional” (i.e., non-disrupting) by SGE were estimated by LASSIE to be under strong negative selection and 9% of “non-functional” mutations were estimated to be under only weak selection. These discordances could in part reflect the influence of natural selection in other cell types or conditions, or limitations of the assay as a measure of disease importance. In any case, they suggest—based on this one gene, cell type, and functional assay—that while there is a strong positive correlation between signatures of natural selection and disease impact, there are nevertheless many exceptions to this general correspondence.

While evolutionary methods clearly have value in predicting Mendelian disease variants, it is less clear that they will be useful for identifying causal variants for complex diseases or other complex traits, many of which segregate at high frequencies, making them unlikely to be under detectable negative selection. Indeed, we found that none of the variant prioritization methods we tested performed well in common variant prediction (Fig. 3e). Interestingly, however, LASSIE and other evolution-based methods performed much better for rare variants associated with complex traits (Fig. 3d), presumably because rare variants tend to have larger effect sizes and experience stronger negative selection. Indeed, recent studies have shown that the effect sizes of GWAS variants are negatively correlated with allele frequencies and allele ages^66, 67^. Together, these observations suggest that evolution-based methods may have an under-appreciated potential for the identification of rare variants associated with complex traits.

We were able to obtain reasonable estimates of allele-specific selection coefficients by pooling data across many genes, thereby “shrinking” estimates toward their average values given the genomic features. As in all such shrinkage strategies, however, the decreased variance in the estimates comes at the cost of increased bias, which will be most evident for genes that are atypical in some way. We attempted to turn this limitation into a strength by using it to reveal classes of genes that showed unusually large or small numbers of variants relative to the LASSIE predictions, corresponding to “enhanced” or “relaxed” strong negative selection. Interestingly, we found that genes under “enhanced” selection are enriched for brain-specific expression and an association with autism spectrum disorder (ASD). This observation is consistent with a reported enrichment for likely gene-disrupting *de novo* mutations in ASD-affected probands relative to unaffected siblings, suggesting the existence of ~400 ASD-associated genes under particularly strong selection^68–71^. Separately, conventional variant-effect predictors have been reported to perform poorly for neurodevelopmental diseases, while gene-level estimates of natural selection—such as pLI and RVIS—perform considerably better^59, 72, 73^, perhaps because the relevant genes are not extraordinarily conserved across species but are under very strong selection in humans. Our findings help to put these observations in an evolutionary context, and suggest that extensions of our methods that unify variant- and gene-level measures of selection could be particularly useful for neurodevelopmental diseases.

### URLs

LASSIE program, https://github.com/CshlSiepelLab/LASSIE/;

LASSIE browser track, http://compgen.cshl.edu/LASSIE/;

UCSC Genome Browser, http://genome.ucsc.edu/;;

dbNSFP database, https://sites.google.com/site/jpopgen/dbNSFP/;

SNVBox database, http://karchinlab.org/apps/appSnvBox.html;

CADD database, https://cadd.gs.washington.edu/;

SIFT database, http://sift.bii.a-star.edu.sg/;

PolyPhen-2 database, http://genetics.bwh.harvard.edu/pph2/;

Eigen database, http://www.columbia.edu/~ii2135/eigen.html;

1000 Genomes project, http://www.internationalgenome.org/;

Roadmap Epigenomics project, http://www.roadmapepigenomics.org/;

SPIDEX database, http://www.openbioinformatics.org/annovar/spidex_download_form.php.

## Acknowledgments

We thank Fernando Racimo and Joshua Schraiber for sharing their source code for inference of selection coefficients, Stephen Wright for providing access to a GPU server at the University of Toronto, Noah Dukler for help with figure preparation, and members of the Siepel laboratory for helpful discussions. This research was supported by US National Institutes of Health grant R35-GM127070. The content is solely the responsibility of the authors and does not necessarily represent the official views of the US National Institutes of Health.

## Online Methods

### Genomic features

For predictive genomic features, we used predefined variant categories indicating the impact of each mutation on the encoded protein, sequence conservation scores, protein structural features, splicing scores, and RNA-seq signals (**Supplementary Table 1**). The variant categories were defined by three indicator variables for whether or not a variant was nonsynonymous, nonsense (stop-gained), or stop-lost in dbNSFP^74^. The conservation scores include scores derived from both protein and multi-species genomic alignments. The protein sequence conservation scores included SIFT^11^, LRT^75^, Mutation Assessor^76^, PROVEAN^77^, SLR^78^, Grantham^79^, PSIC scores from PolyPhen-2 (ref. 21), and HMMEntropy scores from SNV-Box^45^. The nucleotide sequence conservation scores included phyloP scores^8^ derived from vertebrate, mammalian, and primate whole-genome alignments from the UCSC Genome Browser^80^. The protein structural features were obtained from SNV-Box and included predicted secondary structures, and contributions to protein stability, B-factors, and relative solvent accessibilities^45^. We also obtained splicing scores and RNA-seq signals from the non-commercial version of SPIDEX and the Roadmap Epigenomics Project, respectively^81,82^. All features were based on the hg19 (GRCH37) assembly of the human genome.

### Polymorphism data

We obtained 51 high-coverage Yoruba genome sequences from the 1000 Genomes project^1^. To reduce technical errors due to alignment and genotype calling, we applied several filters similar to those used in refs. 33 & 83. These filters eliminated multi-allelic nucleotide sites, simple repeats, transposons, and recent segmental duplicates. Following the same references, we masked all CpG dinucleotides present in either the reference genome or alternative alleles. We also obtained the distributions of ancestral alleles in the human-chimp most recent common ancestor from these same previous studies. In this case, we integrated over these distributions when inferring the global demographic model and mixture model for selection, but then conditioned on the most likely ancestral alelle in the mixture density network for simplicity and efficiency.

### Demographic model and exome-wide distribution of selection coefficients

To obtain sites largely free from selection, we began with the putatively neutral regions defined for INSIGHT^33, 83^. Briefly these regions are defined by excluding all coding exons, conserved noncoding elements, and their close flanking regions. We intersected these regions with the 2-kb flanking regions of all coding exons in the Consensus CDS database^84^ to obtain a subset of putatively neutral sites proximal to coding exons and therefore approximately matched to them in terms of influence from selection from linked sites. We fitted a three-epoch demographic model to the site-frequency spectrum in these exon-proximal “neutral” regions, using Poisson Random Field (PRF) theory for inference (see **Supplementary Note** for details).

We then estimated the bulk distribution of selection coefficients in coding regions under a three-component mixture model, with components for neutral evolution (*s*_0_ = 0), weak negative selection (*s*_1_ < 0), and strong negative selection (*s*_2_ < *s*_1_). This model is defined by the selection coefficients {*s*_0_, *s*_1_, *s*_2_} and three corresponding mixture coefficients, {*w*_0_, *w*_1_, *w*_2_}, where *w_i_* represents the probability that each mutation belongs to component *i* of the model. The free parameters {*s*_1_, *s*_2_, *w*_0_, *w*_1_, *w*_2_} were estimated by maximum likelihood, subject to the constraints that *w*_0_ + *w*_1_ + *w*_2_ = 1, {*w*_0_, *w*_1_, *w*_2_} ≥ 0, with *s*_0_ = 0 held fixed (**Supplementary Note**).

### Training the mixture density model

We trained the mixture density network for inference of allele-specific selection coefficients using mini-batch gradient descent. Data for chromosome 1 was used for testing, data for chromosome 2 was used for validation, and data from the remaining chromosomes was used for training, except for the sex chromosomes (X & Y), which were excluded due to their atypical patterns of mutation and selection, as in previous work. The batch-size was set to 100 nucleotides and the training data were shuffled prior to processing. After each epoch of training, we evaluated the model on the validation set (chromosome 2) and stopped training if the loss (negative log likelihood) did not decrease after five successive epochs (early stopping). Finally, we selected the set of parameters estimated in the epoch with the lowest validation loss. After training, we assigned each potential coding mutation its expected selection coefficient s under the site-and allele-specific distribution defined by the mixture density network, that is, with,

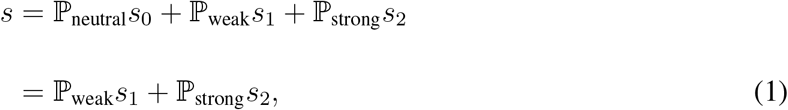

(see Fig. 1c). By construction, *s* ≤ 0, so we generally summarize these estimates using |*s*|, which can be interpreted as a measure of the strength of negative selection. Despite that chromosome X was excluded from the training set, we did generate predictions for this chromosome because it contains many disease variants.

### Comparison with other variant prioritization methods

For comparison with LASSIE, we downloaded precomputed CADD (v1.3; ref. 29), PolyPhen-2 (v2.2.2; ref. 21), SIFT (released in August 2011; ref. 11), Eigen (v1.0; ref. 54), and mammalian phyloP (phyloP46way; ref. 8) scores from their source websites. For all comparisons, we only included variants which were scored by all methods. We visualized the receiver operating characteristic (ROC) curves and calculated the areas under the receiver operating characteristic curve (AUC) using the ROCR package in R^85^.

For the evaluation of Mendelian disease variants, we obtained pathogenic and benign variants from the ClinVar website^55^ in March 2017. Variants annotated as “pathogenic” or “likely pathogenic” were considered “true” disease variants, whereas those annotated as “benign” or “likely benign” were employed as negative controls (ClinVar release in March 2017; ref. 55). Notably, several of the evaluated methods utilized common SNPs or known pathogenic variants in training, which could result in overestimation of their performance. Therefore, we removed all ClinVar variants also in the 1000 Genomes project (phase 3; ref. 1) or the training set of PolyPhen-2. The numbers of positive and negative control variants were balanced by random sampling without replacement. Because true pathological variants are sparse, this matching scheme will tend to result in an over-estimate of the true absolute AUC, but our focus in this article is on the relative performance of the different predictors.

For the evaluation of cancer-driver mutations, we obtained pan-cancer somatic mutations and hotspots of nonsynonymous mutations from ref. 56. We defined cancer genes to be protein-coding genes containing at least one mutational hotspot, cancer-driver mutations to be somatic mutations overlapping mutational hotspots, and passenger mutations to be singleton mutations within cancer genes but not overlapping mutational hotspots. As above, we filtered out all somatic mutations overlapping 1000 Genomes Project variants or the training set of PolyPhen-2, and matched the numbers of cancer diver mutations and passenger mutations by random sampling without replacement.

For the evaluation of nonsynonymous variants associated with complex traits or diseases, we obtained GWAS variants from the GWAS Catalog^58^ (downloaded in November 2017) and identified subsets of rare variants (MAF < 0.01) and common variants (MAF > 0.05). We used nonsynonymous variants from the 1000 Genomes Project as negative controls. After observing that the GWAS variants tended to have higher MAFs than these controls, we matched the distributions of MAFs for the two sets and then randomly sampled negative examples matched in both number and MAF to the GWAS variants, repeating the sampling 100 times to quantify uncertainty.

### Identification of genes under enhanced or relaxed selection

Our model for the expected rates of ultra-rare (MAF < 0.001) variants in the ExAC data^4^ depended critically on an accurate mutation model. Our mutation model was based on precomputed context-dependent mutation rates from the website of the Genome of the Netherlands^60,86^. Because of differences in sample size, we expected that the local mutation rates in the ExAC data would be proportional, but not equal, to the rates estimated from the Genomes of the Netherlands data. Therefore, to recalibrate the local mutation rates, we fitted a simple logistic regression model for the entire genome based on the numbers of ultra-rare synonymous variants in the ExAC data set, assuming that the impact of natural selection on these variants should be minimal. (Note that our finding that few synonymous mutations are under strong negative selection, Fig. 2b, supports this assumption.) Specifically, the logistic regression assumes,

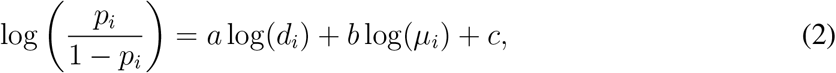

where *p_i_* denotes the probability of observing a rare synonymous mutation *i*, *d_i_* is the sequencing coverage at the corresponding position (which varies considerably in the ExAC data), and *μ_i_* is the corresponding mutation rate estimated by the Genomes of the Netherlands. We validated the mutational model by comparing the observed number of synonymous mutations with the expected number predicted by the mutational model for each gene. We removed short genes (with <200 potential synonymous mutations) and genes for which the mutational model seemed to be misspecified (FDR rate ≤ 0.2) from further consideration.

Based on this recalibrated mutation model, we defined a null model for the number of non-synonymous mutations per gene given the site- and allele-specific selection coefficients estimated by LASSIE. First, we calculated *q_i_*, the expected probability of observing a nonsynonymous mutation *i*, as

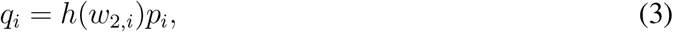

where *p_i_* is the probability of mutation *i* under the neutral mutational model, *w_2,i_* is the probability that mutation *i* is under strong selection as estimated by LASSIE, and h(·) is a mapping from the probabilities of strong selection to relative rates of the nonsynonymous mutation in the ExAC data. To estimate h(·), we grouped all potential nonsynonymous mutations into 1000 equal-width bins based on *w_2,i_* and then estimated h(·) for each selection bin by calculating the ratio of the observed to the expected numbers of nonsynonymous mutations. We then calculated *q_i_* for each nonsynonymous mutation *i* and used *q_i_* to parameterize a Poisson-Binomial distribution describing the null distribution of the number of nonsynonymous mutations for each gene. The probability mass function of the Poisson-Binomial model for a single gene is given by,

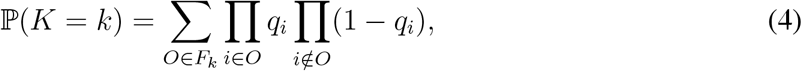

where *k* is the observed number of nonsynonymous mutations in the gene and *F_k_* is the set of all possible arrangements of *k* nonsynonymous mutations. Genes under enhanced or relaxed selections were identified by two separate one-tail tests based on this Poisson-Binomial distribution.

### Gene enrichment analysis

The enrichment analysis of tissue-specific expression was based on annotated tissue-specific genes from the Human Protein Atlas^61^. Only tissues having more than 50 tissue-specific genes were included. The analysis of autism spectrum disorder (ASD) was based on ASD-related genes from the SFARI Gene database^62^. In both cases, significant enrichments were determined using Fisher’s exact test. For the analysis of functional categories, we used PANTHER to investigate the enrichment of both Gene Ontology Slim categories and Reactome pathways^63,87^.

## Code availability

The source code for LASSIE is available on GitHub under the simplified BSD license (http://github.com/CshlSiepelLab/LASSIE/).

## Data availability

The LASSIE scores are available as a UCSC Genome Browser track at http://compgen.cshl.edu/LASSIE/. Additional data generated during the course of our analyses can be obtained from the correspondong author [AS] by request.

## References

1. The 1000 Genomes Project Consortium. A global reference for human genetic variation. Nature 526, 68–74 (2015).

2. The UK10K Consortium. The UK10K project identifies rare variants in health and disease. Nature 526, 82–90 (2015).

3. Mallick, S. et al. The Simons Genome Diversity Project: 300 genomes from 142 diverse populations. Nature 538, 201–206 (2016).

4. Lek, M. et al. Analysis of protein-coding genetic variation in 60,706 humans. Nature 536, 285–291 (2016).

5. Eilbeck, K., Quinlan, A. & Yandell, M. Settling the score: variant prioritization and Mendelian disease. Nat Rev Genet 18, 599–612 (2017).

6. Siepel, A. et al. Evolutionarily conserved elements in vertebrate, insect, worm, and yeast genomes. Genome Research 15, 1034–1050 (2005).

7. Cooper, G. M. et al. Distribution and intensity of constraint in mammalian genomic sequence. Genome Res 15, 901–913 (2005).

8. Pollard, K. S., Hubisz, M. J., Rosenbloom, K. R. & Siepel, A. Detection of nonneutral substitution rates on mammalian phylogenies. Genome Research 20, 110–121 (2010).

9. Gulko, B., Hubisz, M. J., Gronau, I. & Siepel, A. A method for calculating probabilities of fitness consequences for point mutations across the human genome. Nature Genetics 47, 276–283 (2015).

10. Finucane, H. K. et al. Partitioning heritability by functional annotation using genome-wide association summary statistics. Nature Genetics 47, 1228–1235 (2015).

11. Ng, P. C. & Henikoff, S. SIFT: predicting amino acid changes that affect protein function. Nucleic Acids Research 31, 3812–3814 (2003).

12. Cooper, G. M. et al. Single-nucleotide evolutionary constraint scores highlight disease-causing mutations. Nature Methods 7, 250–251 (2010).

13. Lindblad-Toh, K. et al. A high-resolution map of human evolutionary constraint using 29 mammals. Nature 478, 476–482 (2011).

14. Kellogg, E. H., LeaverFay, A. & Baker, D. Role of conformational sampling in computing mutationinduced changes in protein structure and stability. Proteins: Structure, Function, and Bioinformatics 79, 830–838 (2011).

15. Dehouck, Y. et al. Fast and accurate predictions of protein stability changes upon mutations using statistical potentials and neural networks: PoPMuSiC-2.0. Bioinformatics 25, 2537–2543 (2009).

16. Worth, C. L., Preissner, R. & Blundell, T. L. SDM—a server for predicting effects of mutations on protein stability and malfunction. Nucleic Acids Research 39, W215–W222 (2011).

17. Lee, D. et al. A method to predict the impact of regulatory variants from DNA sequence. Nature Genetics 47, 955–961 (2015).

18. Kelley, D. R., Snoek, J. & Rinn, J. Basset: Learning the regulatory code of the accessible genome with deep convolutional neural networks. Genome Research (2016).

19. Zhou, J. & Troyanskaya, O. G. Predicting effects of noncoding variants with deep learning-based sequence model. Nature Methods 12, 931–934 (2015).

20. Alipanahi, B., Delong, A., Weirauch, M. T. & Frey, B. J. Predicting the sequence specificities of dna-and rna-binding proteins by deep learning. Nature Biotechnology 33, 831–838 (2015).

21. Adzhubei, I. A. et al. A method and server for predicting damaging missense mutations. Nature Methods 7, 248–249 (2010).

22. Shihab, H. A. et al. Predicting the functional, molecular, and phenotypic consequences of amino acid substitutions using hidden markov models. Human Mutation 34, 57–65 (2013).

23. Ritchie, G. R. S., Dunham, I., Zeggini, E. & Flicek, P. Functional annotation of noncoding sequence variants. Nature Methods 11, 294–296 (2014).

24. Schwarz, J. M., Cooper, D. N., Schuelke, M. & Seelow, D. MutationTaster2: mutation prediction for the deep-sequencing age. Nature Methods 11, 361– (2014).

25. Jagadeesh, K. A. et al. M-CAP eliminates a majority of variants of uncertain significance in clinical exomes at high sensitivity. Nature Genetics 48, 1581–1586 (2016).

26. Grimm, D. G. et al. The evaluation of tools used to predict the impact of missense variants is hindered by two types of circularity. Human Mutation 36, 513–523 (2015).

27. Huang, Y.-F., Gulko, B. & Siepel, A. Fast, scalable prediction of deleterious noncoding variants from functional and population genomic data. Nature Genetics 49, 618–624 (2017).

28. Gulko, B. & Siepel, A. How much information is provided by human epigenomic data? An evolutionary view. Nature Genetics (2018). In press.

29. Kircher, M. et al. A general framework for estimating the relative pathogenicity of human genetic variants. Nature Genetics 46, 310–315 (2014).

30. Fu, Y. et al. FunSeq2: a framework for prioritizing noncoding regulatory variants in cancer. Genome Biology 15, 480 (2014).

31. di Iulio, J. et al. The human noncoding genome defined by genetic diversity. Nature Genetics 333–337 (2018).

32. Sadri, J., Diallo, A. B. & Blanchette, M. Predicting site-specific human selective pressure using evolutionary signatures. Bioinformatics 27, i266–i274 (2011).

33. Gronau, I., Arbiza, L., Mohammed, J. & Siepel, A. Inference of natural selection from interspersed genomic elements based on polymorphism and divergence. Molecular Biology and Evolution 30, 1159–1171 (2013).

34. Williamson, S. H. et al. Simultaneous inference of selection and population growth from patterns of variation in the human genome. Proceedings of the National Academy of Sciences 102, 7882–7887 (2005).

35. Eyre-Walker, A., Woolfit, M. & Phelps, T. The distribution of fitness effects of new deleterious amino acid mutations in humans. Genetics 173, 891–900 (2006).

36. Keightley, P. D. & Eyre-Walker, A. Joint inference of the distribution of fitness effects of deleterious mutations and population demography based on nucleotide polymorphism frequencies. Genetics 177, 2251–2261 (2007).

37. Boyko, A. R. et al. Assessing the evolutionary impact of amino acid mutations in the human genome. PLoS Genet 4, e1000083 (2008).

38. Kousathanas, A. & Keightley, P. D. A comparison of models to infer the distribution of fitness effects of new mutations. Genetics 193, 1197–1208 (2013).

39. Racimo, F. & Schraiber, J. G. Approximation to the distribution of fitness effects across functional categories in human segregating polymorphisms. PLoS Genet 10, e1004697 (2014).

40. Kim, B. Y., Huber, C. D. & Lohmueller, K. E. Inference of the distribution of selection coefficients for new nonsynonymous mutations using large samples. Genetics 206, 345–361 (2017).

41. Nielsen, R. Molecular signatures of natural selection. Annu. Rev. Genet. 39, 197–218 (2005).

42. Sawyer, S. A. & Hartl, D. L. Population genetics of polymorphism and divergence. Genetics 132, 1161–1176(1992).

43. Evans, S. N., Shvets, Y. & Slatkin, M. Non-equilibrium theory of the allele frequency spectrum. Theoretical Population Biology 71, 109–119 (2007).

44. Bishop, C. M. Mixture density networks. Tech. Rep., Aston University (1994).

45. Wong, W. C. et al. CHASM and SNVBox: toolkit for detecting biologically important single nucleotide mutations in cancer. Bioinformatics 27, 2147–2148 (2011).

46. Zhang, J. & Yang, J.-R. Determinants of the rate of protein sequence evolution. Nature Reviews Genetics 16, 409–420 (2015).

47. Bustamante, C. D., Nielsen, R. & Hartl, D. L. A maximum likelihood method for analyzing pseudogene evolution: implications for silent site evolution in humans and rodents. Mol. Biol. Evol. 19, 110–117 (2002).

48. Kondrashov, F. A., Ogurtsov, A. Y. & Kondrashov, A. S. Selection in favor of nucleotides G and C diversifies evolution rates and levels of polymorphism at mammalian synonymous sites. J. Theor. Biol. 240, 616–626 (2006).

49. Chamary, J. V., Parmley, J. L. & Hurst, L. D. Hearing silence: non-neutral evolution at synonymous sites in mammals. Nature Reviews Genetics 7, 98–108 (2006).

50. Comeron, J. M. Weak selection and recent mutational changes influence polymorphic synonymous mutations in humans. Proceedings of the National Academy of Sciences 103, 6940–6945 (2006).

51. Eory, L., Halligan, D. L. & Keightley, P. D. Distributions of selectively constrained sites and deleterious mutation rates in the hominid and murid genomes. Molecular Biology and Evolution 27, 177–192 (2010).

52. Rasmussen, M. D., Hubisz, M. J., Gronau, I. & Siepel, A. Genome-wide inference of ancestral recombination graphs. PLOS Genetics 10, e1004342– (2014).

53. Yang, Z. Computational Molecular Evolution (Oxford University Press, 2006).

54. Ionita-Laza, I., McCallum, K., Xu, B. & Buxbaum, J. D. A spectral approach integrating functional genomic annotations for coding and noncoding variants. Nature Genetics 48, 214–220 (2016).

55. Landrum, M. J. et al. ClinVar: public archive of relationships among sequence variation and human phenotype. Nucleic Acids Research 42, D980–D985 (2014).

56. Chang, M. T. et al. Identifying recurrent mutations in cancer reveals widespread lineage diversity and mutational specificity. Nat Biotech 34, 155–163 (2016).

57. Findlay, G. M. et al. Accurate classification of BRCA1 variants with saturation genome editing. Nature (2018).

58. MacArthur, J. et al. The new NHGRI-EBI catalog of published genome-wide association studies (GWAS catalog). Nucleic Acids Research 45, D896–D901 (2017).

59. Samocha, K. E. et al. Regional missense constraint improves variant deleteriousness prediction. bioRxiv (2017).

60. Francioli, L. C. et al. Genome-wide patterns and properties of de novo mutations in humans. Nature Genetics 47, 822–826 (2015).

61. Uhlen, M. et al. Tissue-based map of the human proteome. Science 347, – (2015).

62. Abrahams, B. S. et al. SFARI Gene 2.0: a community-driven knowledgebase for the autism spectrum disorders (ASDs). Molecular Autism 4, 36 (2013).

63. Mi, H. et al. PANTHER version 11: expanded annotation data from gene ontology and reactome pathways, and data analysis tool enhancements. Nucleic Acids Research 45, D183–D189 (2017).

64. Cassa, C. A. et al. Estimating the selective effects of heterozygous protein-truncating variants from human exome data. Nature Genetics 49, 806–810 (2017).

65. Keinan, A. & Clark, A. G. Recent explosive human population growth has resulted in an excess of rare genetic variants. Science 336, 740–743 (2012).

66. Zeng, J. et al. Signatures of negative selection in the genetic architecture of human complex traits. Nature Genetics 746–753 (2018).

67. Gazal, S. et al. Linkage disequilibrium-dependent architecture of human complex traits shows action of negative selection. Nature Genetics 49, 1421– (2017).

68. Iossifov, I. et al. The contribution of de novo coding mutations to autism spectrum disorder. Nature 515, 216–221 (2014).

69. O’Roak, B. J. et al. Sporadic autism exomes reveal a highly interconnected protein network of de novo mutations. Nature 485, 246–250 (2012).

70. Sanders, S. J. et al. De novo mutations revealed by whole-exome sequencing are strongly associated with autism. Nature 485, 237–241 (2012).

71. Iossifov, I. et al. De novo gene disruptions in children on the autistic spectrum. Neuron 74, 285–299 (2012).

72. Petrovski, S., Wang, Q., Heinzen, E. L., Allen, A. S. & Goldstein, D. B. Genic intolerance to functional variation and the interpretation of personal genomes. PLOS Genetics 9, e1003709 (2013).

73. Samocha, K. E. et al. A framework for the interpretation of de novo mutation in human disease. Nature Genetics 46, 944–950 (2014).

74. Liu, X., Jian, X. & Eric, B. dbNSFP v2.0: A database of human nonsynonymous snvs and their functional predictions and annotations. Human Mutation 34, E2393–E2402 (2013).

75. Chun, S. & Fay, J. C. Identification of deleterious mutations within three human genomes. Genome Research 19, 1553–1561 (2009).

76. Reva, B., Antipin, Y. & Sander, C. Predicting the functional impact of protein mutations: application to cancer genomics. Nucleic Acids Research 39, e118 (2011).

77. Choi, Y., Sims, G. E., Murphy, S., Miller, J. R. & Chan, A. P. Predicting the functional effect of amino acid substitutions and indels. PLOS ONE 7, 1–13 (2012).

78. Massingham, T. & Goldman, N. Detecting amino acid sites under positive selection and purifying selection. Genetics 169, 1753–1762 (2005).

79. Grantham, R. Amino acid difference formula to help explain protein evolution. Science 185, 862–864 (1974).

80. Casper, J. et al. The UCSC genome browser database: 2018 update. Nucleic Acids Research 46, D762–D769 (2018).

81. Xiong, H. Y. et al. The human splicing code reveals new insights into the genetic determinants of disease. Science 347, 1254806 (2015).

82. Roadmap Epigenomics Consortium et al. Integrative analysis of 111 reference human epigenomes. Nature 518, 317–330 (2015).

83. Arbiza, L. et al. Genome-wide inference of natural selection on human transcription factor binding sites. Nature Genetics 45, 723–729 (2013).

84. Pruitt, K. D. et al. The consensus coding sequence (CCDS) project: Identifying a common protein-coding gene set for the human and mouse genomes. Genome Research 19, 1316–1323 (2009).

85. Sing, T., Sander, O., Beerenwinkel, N. & Lengauer, T. ROCR: visualizing classifier performance in R. Bioinformatics 21, 3940–3941 (2005).

86. Genome of the Netherlands Consortium. Whole-genome sequence variation, population structure and demographic history of the dutch population. Nature Genetics 46, 818–825 (2014).

87. Fabregat, A. et al. The Reactome pathway knowledgebase. Nucleic Acids Research 46, D649–D655 (2018).

